# Ultrasound-Based Hemodynamic Force Trajectories in a Murine Model of Reversible Pressure Overload

**DOI:** 10.64898/2026.01.05.697671

**Authors:** Peyton Day, Thomas Moore-Morris, Craig J. Goergen, Pierre Sicard

**Author notes:** Co-last author Correspondance: Dr. Pierre SICARD, INSERM, CNRS, Université de Montpellier, PHYMEDEXP CHU Arnaud De Villeneuve Bât Crastes de Paulet, 371 Avenue du Doyen Gaston Giraud, 34295 Montpellier cedex, France, Dr. Craig J. Goergen, Weldon School of Biomedical Engineering, Purdue University, West Lafayette, IN, USA.

## Abstract

Hemodynamic forces (HDF) quantify intraventricular pressure gradients as integrated vectors derived from myocardial motion and blood flow, providing a sensitive marker of left ventricular pump efficiency beyond conventional metrics such as ejection fraction or strain. Although HDF analysis has been clinically validated in heart failure, it remains underexplored in preclinical models. To determine whether HDF metrics derived from high-resolution murine echocardiography can sensitively track cardiac dysfunction and predict functional recovery, male mice (n = 8) underwent serial echocardiography at baseline, after transverse aortic constriction (TAC), and four weeks after aortic debanding (deTAC). Left ventricular function and interval-specific HDF components were quantified, including longitudinal and transverse forces over predefined systolic and diastolic windows. Longitudinal interval-based parameters emerged as sensitive, integrative markers of LV dysfunction after TAC and DeTAC. Notably, a novel systolic acceleration-to-relaxation longitudinal HDF ratio showed the strongest associations with pressure overload and subsequent functional recovery (baseline vs TAC: p < 0.001; TAC vs deTAC: p < 0.001; baseline vs deTAC: p = 0.043). In contrast, diastolic HDF indices, including the previously proposed longitudinal e-wave ratio, did not significantly differentiate between time points. Transverse HDF over the systolic impulse interval further demonstrated potential for predicting post-surgical recovery following unloading. These findings support systolic HDF metrics, particularly longitudinal and transverse interval-based parameters, as sensitive integrative markers of LV dysfunction and reverse remodeling in preclinical pressure-overload models, and highlight their translational relevance for echocardiography-based HDF analysis.

**Key points:** Hemodynamic force (HDF) analysis evaluate intraventricular pressure gradients as integrated vectors of myocardial motion and blood flow, providing complementary markers of pump (in)efficiency beyond ejection fraction or strain.

High-resolution murine echocardiography enables robust derivation of interval-specific HDF metrics.

Systolic HDF metrics, particularly transverse components during impulse intervals showed potential for predicting recovery.

Systolic HDF reflect key changes in force generation and intraventricular flow during pressure overload, allowing early detection of maladaptive remodeling and treatment response, with strong translational potential for non-invasive risk stratification in aortic stenosis patients.

## INTRODUCTION

Cardiovascular diseases including heart failure are one of the leading causes of death among people in the developed world, resulting in 19.2 million global deaths in 2023 and impacting 64 million people a year (Stark *et al*., 2025). Despite major therapeutic advances, clinical outcomes remain highly dependent on the ability to accurately monitor disease progression and cardiac recovery, underscoring the need for sensitive and mechanistically meaningful biomarkers of cardiac function (Aimo *et al*., 2022). Conventional imaging metrics, including ventricular volumes, ejection fraction, and myocardial strain, primarily describe myocardial deformation but provide limited insight into the intracavitary forces that govern blood transport. Because ventricular filling and ejection are fundamentally driven by intraventricular pressure gradients (IVPGs), metrics that capture the interaction between myocardial motion and intracardiac fluid dynamics may offer complementary and potentially earlier indicators of functional impairment and recovery (Sørensen *et al*., 2025). In this context, hemodynamic forces (HDF) have emerged as quantitative descriptors of cardiac performance by estimating the pressure forces responsible for blood acceleration within the heart (Vallelonga *et al*., 2021). HDF reflect the dynamic balance between blood inertia and pressure fields generated by myocardial contraction and relaxation, thereby providing a direct mechanical link between myocardial deformation, chamber geometry, and intracardiac flow dynamics (Greenberg *et al*., 2001). Initially introduced more than two decades ago using Doppler M-mode echocardiography (Greenberg *et al*., 2001), HDF analysis was subsequently validated using 4D Flow MRI (Pedrizzetti *et al*., 2014, 2017) and has demonstrated clinical relevance in several pathological settings. Notably, recent studies have shown that HDF metrics can predict long-term responses to cardiac resynchronization therapy in patients with left bundle branch block (Eriksson *et al*., 2017; Laenens *et al*., 2024), highlighting their potential as clinically actionable biomarkers of ventricular mechanics. Despite these promising clinical results, HDF metrics remain largely underexplored in preclinical research, particularly in small animal studies. Although, one study employed MRI-based approaches for HDF assessment (Daal *et al*., 2022), no investigations to date have leveraged high-resolution echocardiography in rodent models, an advancement that could enable cost-effective longitudinal studies with greater accessibility and earlier detection of pathologies.

The present study addresses this gap by exploring a novel approach to HDF quantification using high-resolution ultrasound imaging in a murine model of reverse pressure overload. The TAC/deTAC model enables the evaluation of dynamic changes in ventricular mechanics during loss and recovery of function. Given the inherent complexity of HDF waveforms and their derived parameters based on intraventricular pressure gradients, a central aim of this work is to identify HDF metrics that are most representative of cardiac functional status and to establish robust quantification strategies for their extraction in a preclinical ultrasound setting.

## MATERIALS AND METHODS

### Transverse Aortic Constriction (TAC) and Aortic Debanding (deTAC)

Experiments were conducted following European Directive (2010/63/EU) and French laws for laboratory animal use. The local institutional ethics committee (CCEA n°36) approved this study (no. 2023013113485359). We used C57BL/6JRj male mice aged 8-week-old (n=8); As previously described, we induce heart failure using transverse aortic constriction (TAC) (Ghajar-Rahimi *et al*., 2025). Following the recovery period, we injected the mice twice a day with buprenorphine (1mg/kg, s.c.) for further pain control. Minimally invasive aortic debanding (deTAC) procedure was performed when EF was measured below 45% (Ghajar-Rahimi *et al*., 2025) and we injected the mice twice a day with buprenorphine (1mg/kg, s.c.) for further pain control. We followed the ARRIVE guidelines 2.0 for animal reporting (Percie du Sert *et al*., 2020)

#### Ultrasound Imaging

Vevo 3100 high-frequency ultrasound system (FUJIFILM VisualSonics) with a 40 MHz center frequency MX550D linear array probe was used to assess cardiac function on mice under isoflurane anesthesia, as previously described (Ghajar-Rahimi *et al*., 2025). We performed two-dimensional (2D) echocardiography following American Physiological Society guidelines for assessment of murine LV cardiac function (Lindsey *et al*., 2018). The core temperature was maintained between 36-37.5°C, and the heart rate was monitored and maintained >450 beats/minute throughout the procedure. We used the parasternal long-axis view to collect B-mode images of the left ventricle long axis view, parasternal short axis view at a frame rate of 250 frames/sec. We collected four-dimensional (4D) LV images in short-axis using a linear step motor (step size= 0.13 mm) to scan from the base of the apical epicardium to the ascending aorta to measure the mitral valve diameter at its maximal opening during diastole. If 4D ultrasound was unavailable we used a 2D short-axis view collected at the LV base. Aortic valve diameter was measured from the PLAX view between the leaflets at their maximal systolic opening. All valve diameters were measured perpendicular to the direction of blood flow.

#### Data Processing and Cycle Definition

Offline images were used to calculate 2D LV strain and HDF analysis using Vevo Strain Software 2.0 (FUJIFILM VisualSonics), from the 2D parasternal long-axis (PLAX) B-mode images.

Endocardial and myocardial boundaries were delineated at end-systole and end-diastole using the Vevo Strain software, with speckle-tracking used to calculate intermittent boundary traces. The software-placed end-systole and end-diastole markers were manually verified and, if necessary, corrected to align with the absolute maximum and minimum LV volumes, respectively.

The cardiac cycle was defined from the resultant LV volume curve. Systole was defined as the interval from the point of maximum LV volume (end-diastole) to the first local minimum LV volume (end-systole). Diastole comprises the interval from end-systole to the subsequent end-diastole.

#### HDF Component and Interval Definitions

HDF was resolved into longitudinal (apical-basal) and transverse (lateral-septal) components, assuming radial symmetry. The z-axis was defined originating at the LV apex (z=0) and extending through the basal tissue between the mitral and aortic valves. Specific diastolic and systolic intervals were defined from the HDF curve and when possible, Root Mean Square (RMS) values of HDF metrics were calculated over their respective intervals to improve measurement accuracy. These include the following:

**Diastolic E-Wave:** Defined as the interval during diastole commencing at the first positive-going zero-crossing of the HDF curve and concluding at the subsequent zero-crossing. If the A-wave did not produce a negative HDF value, the E-wave was defined as ending at the inflection point between the E and A waves. If no A-wave was present, the E-wave extended to the end of diastole.
**Systolic Intervals:** The systolic peak was identified as the maximal HDF value during systole.
**Systolic Impulse:** The total interval from the start of systole (or first positive HDF value) until the HDF zero-crossing immediately preceding end-systole.
**Systolic Acceleration:** The interval from the HDF zero-crossing (or systolic minimum) immediately preceding the systolic peak, up to the systolic peak.
**Systolic Relaxation:** The interval from the systolic peak to the end of the systolic impulse (the zero-crossing immediately preceding end-systole).

## Statistical Analysis

All statistical analysis was performed with R version 4.5.1 (RStudio), and the data are presented as mean ± standard deviation (SD). We used a one-way repeated measures ANOVA with Tukey post-hoc multiple comparisons test to compare baseline, 3-week TAC, and 4-week deTAC global parameters, checking for normality with a Shapiro-Wilk test. A Kruskal-Wallis test with Dunn’s multiple comparisons test was used for non-normally distributed data. For interval timing metrics analysis, we used both the raw and normalized data for each timing metric to the overall RR interval length. We used a Pearson correlation to describe relationships between 3-weeks TAC and 4-weeks post-deTAC variables.

Principal component analysis (PCA) was run to determine new meaningful underlying variables, and included all cardiovascular metrics evaluated with high-resolution ultrasound. We used a standardized PCA method due to the different measurement scales of the variables included. Principal components (PCs) were selected based on parallel analysis with 1000 simulations and a 95% percentile level. We plotted the first two PCs against one another to distinguish between the two groups. The contributions of a variable to PC1 and PC2 were averaged to evaluate the overall importance. We set the threshold for statistical significance at *p*<0.05.

## RESULTS

### Left Ventricular Systolic Hemodynamic Performance Recovers in Accordance with Classical Parameters

High-resolution ultrasound was used to assess LV anatomical and functional trajectories at baseline, during TAC, and 4 weeks after deTAC (Figures 1A-C and Supplemental Table 1). As previously described (Ghajar-Rahimi *et al*., 2025; Salvas *et al*., 2025*a*), LV ejection fraction and LV longitudinal strain decreased during TAC and recovered after debanding (Supplemental Table 1). To calculate HDF, we measured the mitral and aortic valves. Using 4D short-axis scans at the base of the heart, we obtained a clear view of the mitral valve (Figure 1A), and a 2D long-axis view was used to measure the aortic valve. While aortic valve area increased under TAC conditions before returning to baseline following debanding, mitral valve area remained stable (Supplemental Table 1). Using VevoStrain software 2.0, we reconstructed full-cycle HDF metrics, including transverse and longitudinal HDF (Figure 1B), along with polar plots to assess HDF distribution in the left ventricle (Figure 1D).

**Figure 1.**
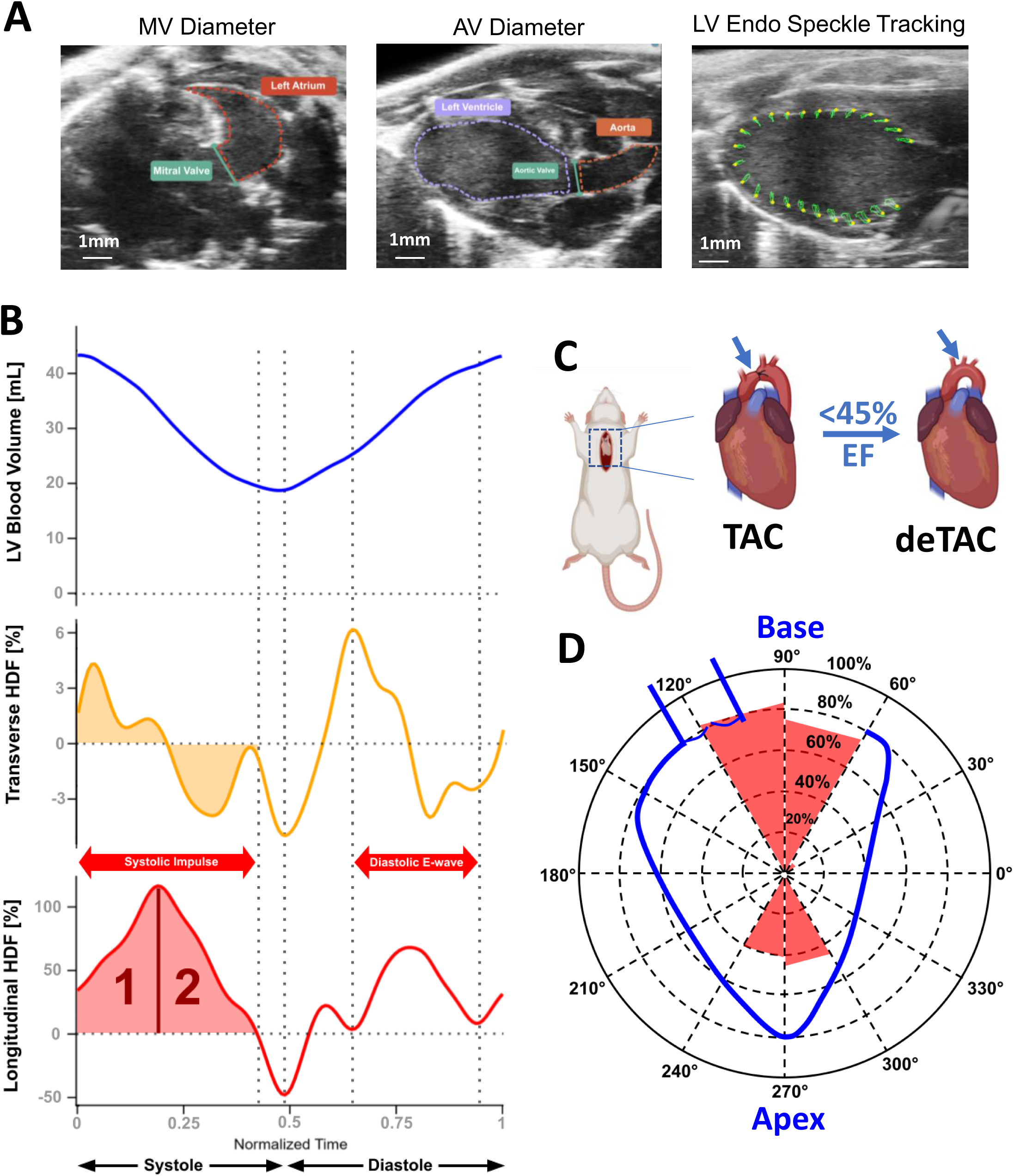
**Echocardiographic Characterization of Left Ventricular Hemodynamic Force (HDF) in a Murine Model**. (A) labeled images of the three ultrasound scans used to determine the inputs for HDF calculation. From left to right, a short axis (SAX) view of the atrium for determining mitral valve (MV) diameter, a parasternal long axis (PLAX) view of the left ventricle to determine aortic valve (AV) diameter, and another PLAX view of the left ventricle recorded over the cardiac cycle to conduct speckle tracking. (B) Representative curves of typical left ventricular (LV) volume (top, blue) and the corresponding transverse (middle, orange) and longitudinal (bottom, red) components of hemodynamic force plotted over a single, time-normalized cardiac cycle. (C) Visual summary of the transverse aortic constriction (TAC) procedure followed by the debanding (deTAC) procedure. (D) Typical polar plot output from HDF calculations overlayed with an anatomical outline of the LV in blue labeled with relevant reference points.

Analysis of the overall longitudinal HDF profile (Figures 1B; Figure 2A-B) shows that, under baseline conditions, forces rapidly rise at the onset of systole to a positive peak, followed by a gradual decline to a minimum marking the transition to diastole. During diastole, longitudinal forces increased again with minor fluctuations, displaying a distinct E-wave peak and a final decrease during late diastolic filling (A-wave). To quantify overall force amplitude, we compared the RMS values of longitudinal HDF across the entire R-R interval between baseline, TAC, and deTAC conditions (Figure 2C). Unexpectedly, no significant difference was observed between baseline and TAC. In contrast, deTAC hearts exhibited a significantly higher longitudinal HDF compared to TAC (*p*<0.05). We next assessed which phase of the cardiac cycle, systole or diastole, provides the most informative parameters to discriminate longitudinal HDF alterations (Figures 2D–E; Table 1). The systolic impulse was significantly reduced in TAC hearts compared to both baseline and deTAC (*p*<0.001, Figure 2D), whereas diastolic longitudinal HDF showed no significant differences among groups (Figure 2E).

**Figure 2.**
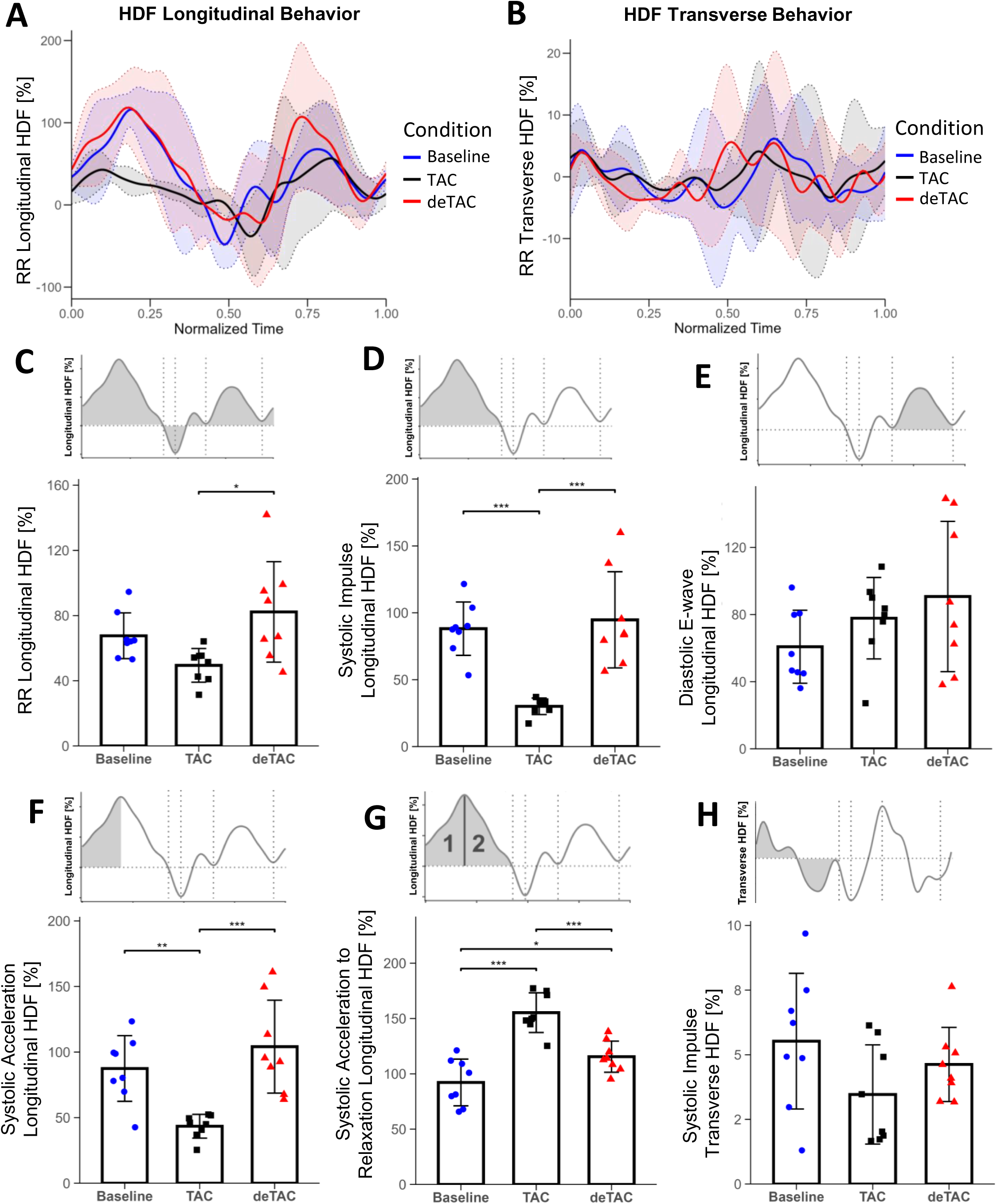
Phase-specific and derived analysis of hemodynamic force (HDF) parameters in TAC/deTAC model. (A) Mean longitudinal HDF behavior over a normalized RR interval for baseline, TAC, and deTAC conditions ± SD. (B) Mean transverse HDF behavior over a normalized RR interval for baseline, TAC, and deTAC conditions ± SD. (C) Longitudinal HDF across the entire RR interval; longitudinal baseline condition in blue, TAC condition in gray, and deTAC condition in red. The trace above each graph represents the baseline mean graph behavior with the region of interest highlighted. (D) Longitudinal HDF isolated to the systolic impulse phase, representing the primary force generation section of systole. (E) Diastolic E-wave longitudinal HDF, representing the primary region of interest in the diastolic phase for HDF. (F) The systolic acceleration component of longitudinal HDF, representing a more specific interval of systolic force generation. (G) A novel ratio of systolic acceleration to relaxation HDF, calculated by dividing the systolic acceleration (area labeled 1) by the systolic relaxation (area labeled 2). (H) Transverse HDF isolated to the systolic impulse phase, representing the vortex forming components in the left ventricle during systolic impulse. *Statistical notations: *p < 0.05, **p < 0.01, ***p < 0.001*.

**Table 1.** HDF cardiac parameter intergroup comparisons in baseline, TAC, and deTAC Groups.

|  |  |  |  |  |  |  |  |
| --- | --- | --- | --- | --- | --- | --- | --- |
| Metric | Method | Baseline (Mean ± SD) | TAC (Mean ± SD) | deTAC (Mean ± SD) | Baseline vs TAC (p) | Baseline vs deTAC (p) | TAC vs deTAC (p) |
| RR Length | Kruskal-Wallis | 120.88 ± 9.28 | 113.00 ± 13.98 | 116.00 ± 15.81 | 0.204 | 0.482 | 0.405 |
| RR Longitudinal RMS | ANOVA | 67.60 ± 14.04 | 49.44 ± 10.36 | 82.24 ± 30.82 | 0.202 | 0.343 | 0.011 (*) |
| RR Transverse RMS | ANOVA | 7.04 ± 2.84 | 6.88 ± 3.01 | 7.05 ± 4.01 | 0.995 | 1 | 0.994 |
| RR Ratio RMS | Kruskal-Wallis | 10.55 ± 3.82 | 14.21 ± 6.44 | 8.53 ± 3.73 | 0.353 | 0.512 | 0.197 |
| RR Angle | ANOVA | 80.50 ± 3.38 | 78.75 ± 3.88 | 82.00 ± 3.02 | 0.576 | 0.664 | 0.168 |
| Systole Length | ANOVA | 53.88 ± 6.29 | 58.38 ± 10.06 | 54.62 ± 15.75 | 0.713 | 0.99 | 0.789 |
| Systolic Impulse Length | ANOVA | 43.12 ± 8.03 | 50.75 ± 9.97 | 45.62 ± 10.20 | 0.262 | 0.858 | 0.534 |
| Systolic Acceleration Length | ANOVA | 16.88 ± 5.41 | 9.12 ± 2.42 | 20.75 ± 6.69 | 0.018 (*) | 0.31 | <0.001 (***) |
| Systolic Relaxation Length | ANOVA | 24.12 ± 5.96 | 39.12 ± 7.59 | 23.25 ± 5.28 | <0.001 (***) | 0.959 | <0.001 (***) |
| Systole Length Normalized | ANOVA | 44.68 ± 5.09 | 52.12 ± 10.02 | 47.09 ± 11.26 | 0.26 | 0.861 | 0.527 |
| Systolic Impulse Length Normalized | Kruskal-Wallis | 35.85 ± 7.09 | 45.32 ± 9.76 | 39.60 ± 8.10 | 0.121 | 0.305 | 0.458 |
| Systolic Acceleration Length Normalized | ANOVA | 14.06 ± 4.49 | 8.18 ± 2.46 | 17.83 ± 5.20 | 0.028 (*) | 0.197 | <0.001 (***) |
| Systolic Relaxation Length Normalized | Kruskal-Wallis | 19.96 ± 4.78 | 34.85 ± 7.01 | 20.51 ± 6.56 | 0.004 (**) | 0.75 | 0.003 (**) |
| Systolic Acceleration to Relaxation Length Ratio | ANOVA | 76.90 ± 40.24 | 24.39 ± 9.05 | 94.98 ± 39.82 | 0.012 (*) | 0.529 | <0.001 (***) |
| Systolic Impulse Longitudinal RMS | ANOVA | 88.15 ± 19.95 | 30.10 ± 6.15 | 94.75 ± 35.95 | <0.001 (***) | 0.848 | <0.001 (***) |
| Systolic Impulse Transverse RMS | Kruskal-Wallis | 5.53 ± 2.62 | 3.46 ± 1.92 | 4.62 ± 1.43 | 0.268 | 0.524 | 0.432 |
| Systolic Impulse AUC | Kruskal-Wallis | 31.91 ± 11.83 | 4.70 ± 2.04 | 31.35 ± 22.20 | <0.001 (***) | 0.572 | 0.003 (**) |
| Systolic Impulse Angle | ANOVA | 83.75 ± 3.11 | 81.25 ± 5.99 | 85.38 ± 1.06 | 0.429 | 0.693 | 0.116 |
| Systolic Peak | Kruskal-Wallis | 129.17 ± 28.89 | 55.92 ± 10.03 | 141.51 ± 58.55 | 0.002 (**) | 0.944 | 0.001 (**) |
| Systolic Acceleration Longitudinal RMS | ANOVA | 87.47 ± 25.04 | 43.45 ± 9.05 | 104.10 ± 35.42 | 0.007 (**) | 0.411 | <0.001 (***) |
| Systolic Relaxation Longitudinal RMS | Kruskal-Wallis | 95.98 ± 26.72 | 28.02 ± 5.34 | 91.76 ± 37.38 | <0.001 (***) | 0.572 | 0.003 (**) |
| Late Systole Longitudinal RMS | Kruskal-Wallis | 41.25 ± 15.55 | 19.96 ± 19.94 | 37.67 ± 11.86 | 0.065 | 0.805 | 0.06 |
| Systolic Acceleration to Relaxation Longitudinal Ratio | ANOVA | 92.22 ± 21.12 | 155.31 ± 17.95 | 115.50 ± 14.09 | <0.001 (***) | 0.043 (*) | <0.001 (***) |
| E-wave Longitudinal RMS | ANOVA | 60.81 ± 21.77 | 77.81 ± 24.27 | 90.74 ± 44.80 | 0.547 | 0.172 | 0.702 |
| E-wave Ratio RMS | ANOVA | 12.24 ± 8.02 | 12.74 ± 6.32 | 9.10 ± 7.30 | 0.99 | 0.667 | 0.583 |
| E-wave Angle | ANOVA | 81.75 ± 4.46 | 79.62 ± 4.41 | 82.00 ± 6.05 | 0.68 | 0.995 | 0.619 |

Statistical notations: \**p* < 0.05, \*\**p* < 0.01, \*\*\**p* < 0.001.

Given these findings, we focused on a detailed analysis of systolic dynamics by calculating more than fifteen systolic force parameters (Table 1). Among these parameters, the systolic acceleration RMS of longitudinal HDF was significantly reduced in TAC hearts compared to both baseline and deTAC hearts (*p*<0.01 and *p*<0.001, Figure 2F)). Interestingly, the systolic acceleration-to-relaxation ratio was significantly increased during TAC compared to baseline and deTAC (*p*<0.001, Figure 2G). Notably, this ratio did not return to baseline values after deTAC and remained significantly elevated relative to baseline (*p*<0.05). Finally, analysis of systolic transverse impulse parameters revealed no significant differences between groups (Figure 2B and H, Table 1).

### Interval-Specific Hemodynamic Forces Improve Prediction of Ventricular Remodeling Compared to Strain

We then assessed the degree to which HDF metrics correlate with classical cardiovascular function, such as ejection fraction, LV strain and aortic peak velocity, by conducting Pearson correlation tests. The overall correlation behavior of metrics of interest are summarized in the inter-correlation matrix (Supplemental Figure 1) which identifies two distinct sets of groups in the clustered blue and red regions (Figure 3A). Most HDF metrics show a positive correlation with LV ejection fraction and aortic peak velocity, except for the e-wave longitudinal RMS, which behaves unpredictably, and the systolic acceleration–to–relaxation ratio, which shows a predictable but inverse relationship (Figure 3A). Interestingly, transverse component of the systolic impulse, which was not significant in intergroup comparisons was found to be significant across all groups (r=0.42, *p*=0.043, Supplemental Table 2) and the paired TAC metric to deTAC LV ejection fraction (r=-0.82, *p* =0.012, Supplemental Table 3) when conducting Pearson correlation tests (Figure 3D).

**Figure 3.**
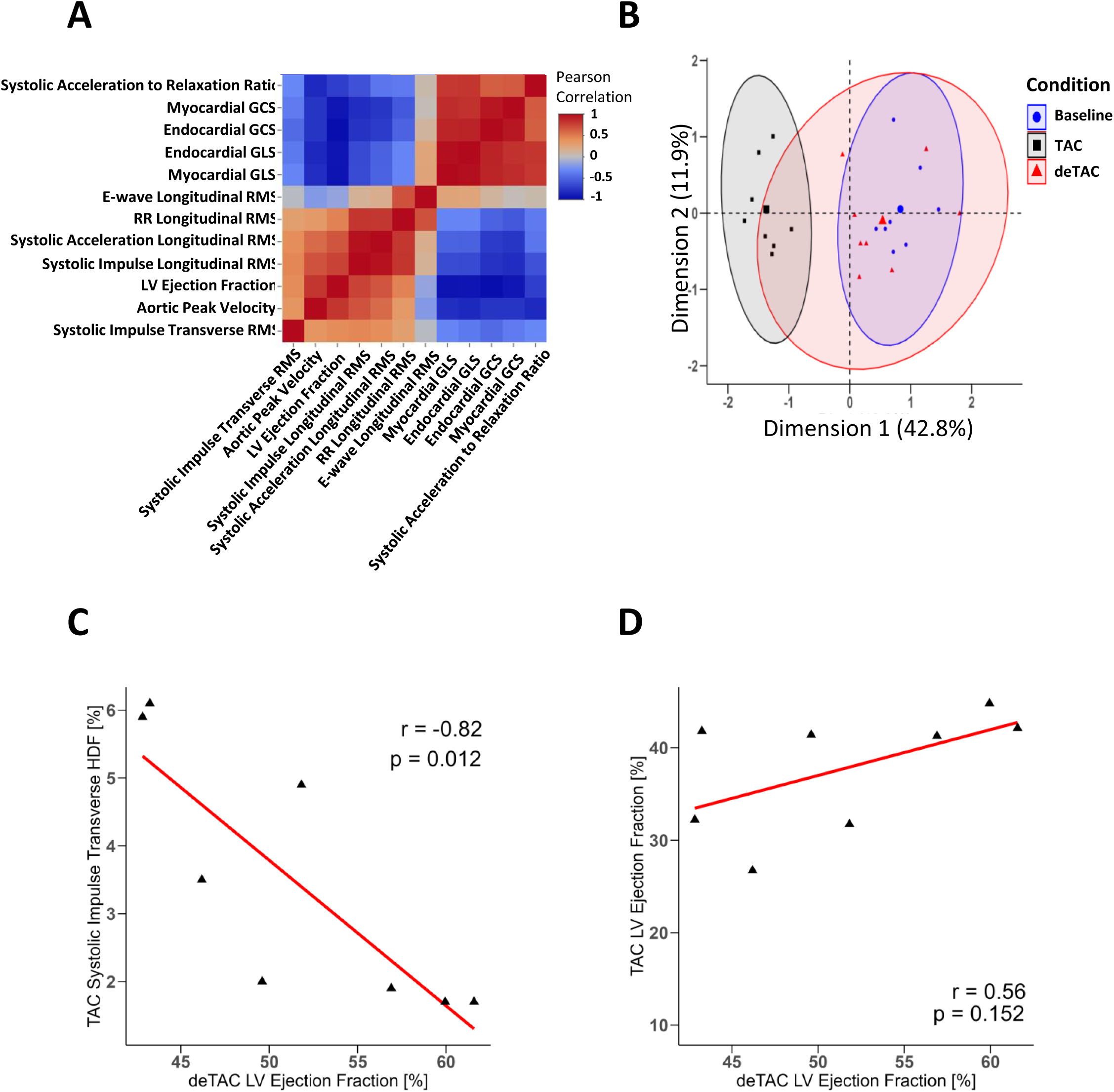
Correlative analysis of HDF and traditional echocardiographic parameters. (A) Inter-correlation matrix of strain, ejection fraction, and aortic peak velocity against various HDF parameters. The color scale indicates the strength and direction of the correlation, where deep red signifies a strong positive correlation (r ≈ 1) and deep blue signifies a strong negative correlation (r ≈ -1). The distinct color blocks demonstrate that strain parameters and the derived HDF ratio metric are highly correlated with one another as well as that ejection fraction and aortic peak velocity are highly correlated with longitudinal HDF parameters. (B) Partial Component Analysis (PCA) of individual mouse samples across all metrics measured. The high variability of the deTAC condition demonstrates how broader and many diastolic HDF parameters do not show recovery after deTAC. (C) Grouped correlation of TAC condition systolic impulse transverse HDF against deTAC LV ejection fraction recovery (r=-0.82, p=0.012). (D) Grouped correlation of TAC condition LV ejection fraction against deTAC LV ejection fraction recovery (r=0.56, p=0.152).

PCA was then applied to test whether combined HDF and functional metrics could discriminate between experimental conditions. The first two principal components explained 55% of the total variance (PC1: 43%, PC2: 12%; Supplemental Figure 2 A), with baseline and TAC groups showing partial separation along PC1 and deTAC animals occupying an intermediate position, consistent with incomplete reversal of the TAC-induced alterations (Figure 3B). PC1 was mainly driven by aortic peak velocity, endocardial GLS, LV ejection fraction, and systolic strain indices, and therefore reflected a global systolic function/loading axis with TAC hearts at one extreme, baseline at the opposite, and deTAC in between. In contrast, PC2 was dominated by transverse HDF amplitude (cycle_trans_rms), systolic longitudinal HDF length (sys_im_length_norm), e-wave longitudinal HDF (e_wave_long_rms), and cycle length (cycle_length). This captured a hemodynamic force pattern/timing axis that separated animals by transverse force magnitude, spatial extent, and cycle duration, not conventional systolic indices (Supplemental Figure 2A-C). TAC induced a distinct multivariate HDF signature, only partially normalized after debanding and missed by univariate parameters. Finally, we assessed whether any HDF or conventional metrics measured during TAC could predict the reversal of LV functional impairment after debanding. Using Pearson correlation analysis (Supplemental Tables 2 and 3), we found that among the 41 parameters tested, only HDF-derived metrics, specifically the systolic transverse RMS and systolic transverse impulse, significantly correlate with reverse LV functional recovery (Figure 3C), whereas ejection fraction failed to show any predictive value (Figure 3D).

## DISCUSSION

Our preclinical study demonstrates that HDF parameters derived from high-resolution ultrasound reliably track cardiac function and closely align with established markers such as global longitudinal strain, circumferential strain, and left ventricular ejection fraction. Beyond this concordance, our findings indicate that HDF metrics offer added value by improving the prediction of left ventricular remodeling and functional recovery following surgical intervention. Together, these results highlight the potential of HDF analysis to enhance prognostic assessment of LV recovery beyond conventional echocardiographic measures.

Recent advances in ultrasound and cardiac imaging analysis methodologies have rapidly expanded the tools available to clinicians and researchers, enabling deeper, faster, and more accurate assessment of cardiac function during disease progression (Gillam & Marcoff, 2024). Beyond conventional measures such as ejection fraction, segmental cardiac strain has markedly improved the early detection of developing heart failure following myocardial infarction or pressure overload, including aortic constriction, by revealing subtle patterns of myocardial deformation abnormalities (Smiseth *et al*., 2025). Initially developed for clinical applications, cardiac strain analysis has now been incorporated into recent guidelines for assessing cardiac function in rodent models (Lindsey *et al*., 2018; Zacchigna *et al*., 2020). Because LV function cannot be fully captured by ejection fraction alone, the heart must be considered as an integrated system in which cavity interdependence plays a critical role. Consequently, holistic approaches could be used to better understand cardiac dysfunction trajectory, including the assessment of left atrial functional dynamics (Salvas *et al*., 2025*b*). More recently, the interaction between myocardial motion and intracardiac fluid dynamics, assessed by cardiac MRI and increasingly by advanced ultrasound techniques, has gained attention. Early evaluation of HDF has been shown to carry prognostic value in asymptomatic patients and may provide unique insights into myocardial dysfunction that extend beyond traditional echocardiographic parameters and deformation-based measures (Vallelonga *et al*., 2021). Unlike global strain or LVEF, which summarize deformation or volume change, systolic HDF evaluate the magnitude and orientation of intraventricular pressure gradients, so early maladaptive remodeling appears as “misaligned” or more transverse forces even when strain and EF are still within preserved or only mildly reduced ranges (Backhaus *et al*., 2022). Specifically, HDF analysis has been shown to differentiate patients who recover LV function after transcatheter aortic valve implantation (TAVI) from those with persistent hemodynamic dysfunction (Vairo *et al*., 2023; Angotti *et al*., 2025). In this preclinical study, we successfully applied ultrasound-based HDF analysis in a mouse model of aortic banding and debanding that mimics the treatment of aortic stenosis. Following debanding, systolic HDF components and total left ventricular longitudinal force (LVLF) improved significantly, indicating that recovery of contractile function is accompanied by a marked normalization of intraventricular fluid dynamics (Vairo *et al*., 2023; Angotti *et al*., 2025). While strain and ejection fraction metrics correlate well with aortic peak velocity, an established indicator of LV systolic performance, our findings suggest that, compared with systolic HDF components, these conventional metrics have less predictive power when used as single time-point measurements to anticipate functional recovery after surgical intervention.

We tested for any specific variation in effects for diastolic and systolic regions of interest. In particular, previous reports have found that the regions of interest are the systolic impulse (Faganello *et al*., 2024; Li *et al*., 2025) and diastolic e-wave regions (Eriksson *et al*., 2016; Backhaus *et al*., 2022). Generally, in our TAC and deTAC model systolic impulse metrics were found to have much higher significance as compared to the diastolic e-wave metrics. Further, the more specified the intervals became, the more significant that the intergroup statistical power became. These findings highlight the particular value of temporally and spatially resolved systolic HDF descriptors in characterizing disease severity and reversibility in this model. Consistent with recent 4D flow phase contrast MRI–based studies indicating that the prognostic value of current HDF implementations in HFpEF depends strongly on parameter selection and analytical context, our data support the need for careful metric choice and contextual interpretation to extract robust prognostic information from HDF analysis (Arvidsson *et al*., 2022). Building on this, PCA of combined HDF and conventional echocardiographic variables revealed that systolic HDF features contributed most strongly to the main axis separating baseline, TAC, and deTAC conditions, whereas diastolic e-wave related metrics had only modest influence on group discrimination. The data thus reflects the high variability observed during diastole, as seen in both longitudinal and transverse HDF components. This suggests that, at least in the context of pressure overload and its relief, altered systolic force generation and redirection within the ventricle capture the dominant mechanisms driving maladaptive remodeling, while diastolic HDF changes may be subtler, secondary phenomena. Focusing on carefully defined systolic windows therefore appears to enhance both the sensitivity and the statistical power of HDF-based analyses, supporting their use as potential early integrative markers of LV contractile recovery after unloading interventions (Vallelonga *et al*., 2021; Vairo *et al*., 2023; Angotti *et al*., 2025). Taken together, these results suggest that echocardiographic assessment of HDFs may provide a powerful tool to distinguish responders from non-responders after surgical treatment, thereby identifying individuals at risk of persistent hemodynamic dysfunction.

### Limitation

The use of 2D speckle-tracking echocardiography in Vevo Strain 2.0 for HDF calculations relies on assumptions from a single PLAX view, which cannot capture true 3D myocardial motion including out-of-plane rotation and deformation, leading to underestimation of transverse strain components; transitioning to 4D HDF analysis could enhance reproducibility by enabling full volumetric tracking from a single dataset, while potentially revealing finer details such as diastolic e-wave parameters and angle variations that exhibit high variability in 2D assessments.

Additionally, our analysis emphasized systolic HDF metrics due to their superior statistical power and predictive value in this TAC/deTAC model, potentially overlooking diastolic e-wave contributions that could offer complementary insights into filling dynamics and vary across different heart failure etiologies. While future work will be needed to create analysis strategies that fully combine a wider variety of metrics to fully characterize cardiac remodeling.

## CONCLUSION

This preclinical study establishes ultrasound-derived HDF metrics, particularly systolic transverse impulse components, as possible predictors of LV functional recovery following aortic debanding in a mouse TAC model, outperforming conventional strain and ejection fraction measures at single time points. These findings underscore the value of integrating intraventricular fluid dynamics with myocardial deformation for early identification of responders versus non-responders to unloading interventions. While future investigations incorporating 3D/4D HDF analysis, diastolic-focused assessments, and sex-balanced cohorts could refine their clinical translatability in pressure overload heart failure, this work demonstrates the potential of HDF waveform analysis to improve assessment of LV remodeling over standard echocardiographic measures.

## Data Availability Statement

The data that support the findings of this study are available from the corresponding author upon reasonable request.

## Author Contributions

**Peyton Day:** Formal Analysis, Investigation, Visualization, Writing – original draft, review & editing. **Thomas Moore-Morris:** Conceptualization, Investigation, Methodology, Supervision, Writing – original draft, review & editing. **Craig J Goergen:** Conceptualization, Methodology, Supervision, Writing – review & editing. **Pierre Sicard:** Conceptualization, Investigation, Methodology, Supervision, Writing – original draft, review & editing.

## Acknowledgements

We gratefully thank the staff for animal housing (PhyMedExp). We acknowledge the technical support provided by Imagerie du Petit Animal de Montpellier (IPAM) for accessing high-resolution ultrasound (LRQA Iso9001; France Life Imaging (grant ANR-11-INBS-0006); IBISA; Leducq Foundation (RETP), I-Site Muse).

---

**Supplemental Figure 1.**
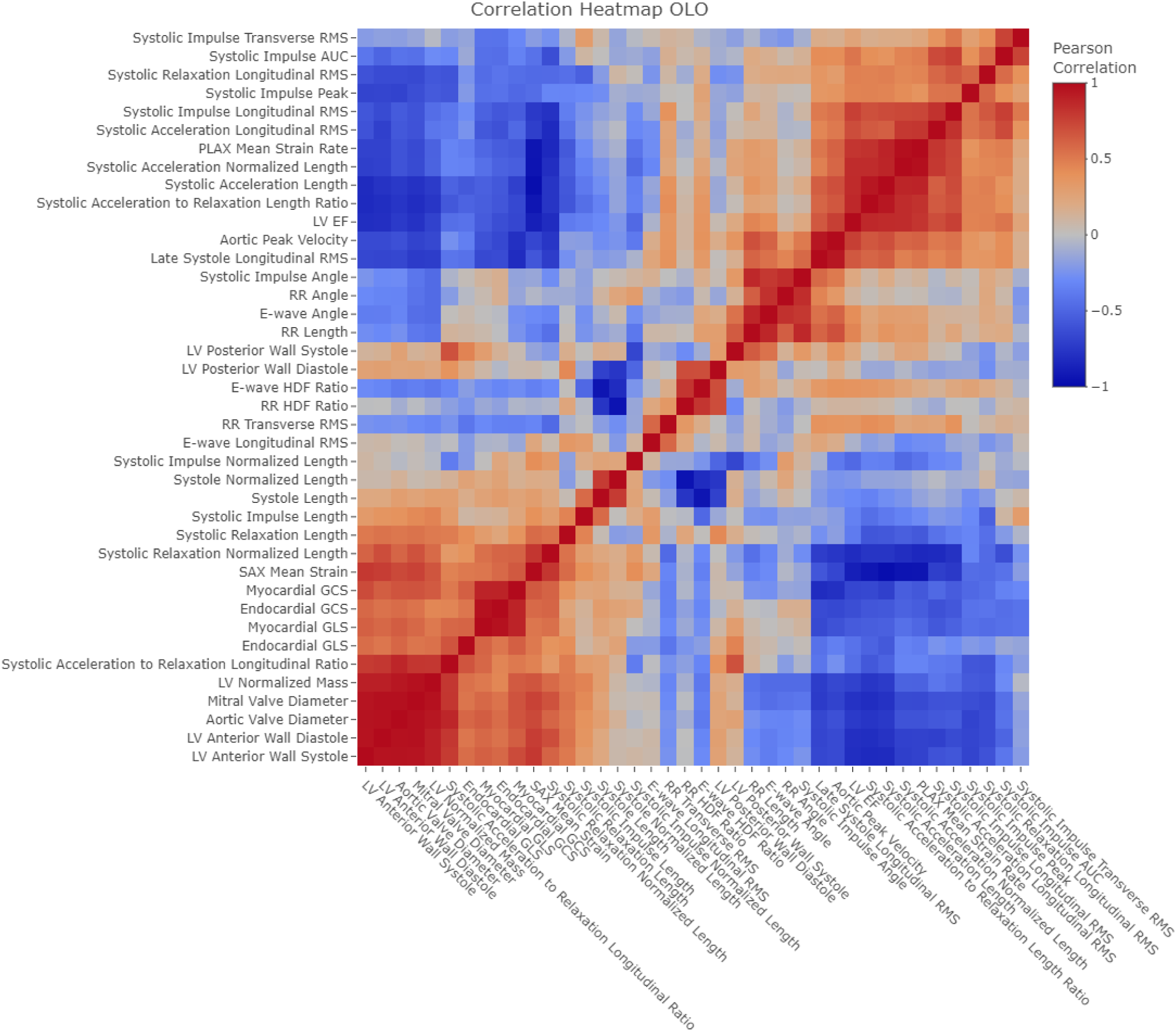
Overall correlation behavior of metrics of interest are summarized in the inter-correlation matrix The color scale indicates the strength and direction of the correlation, where deep red signifies a strong positive correlation (r ≈ 1) and deep blue signifies a strong negative correlation (r ≈ -1).

**Supplemental Figure 2.**
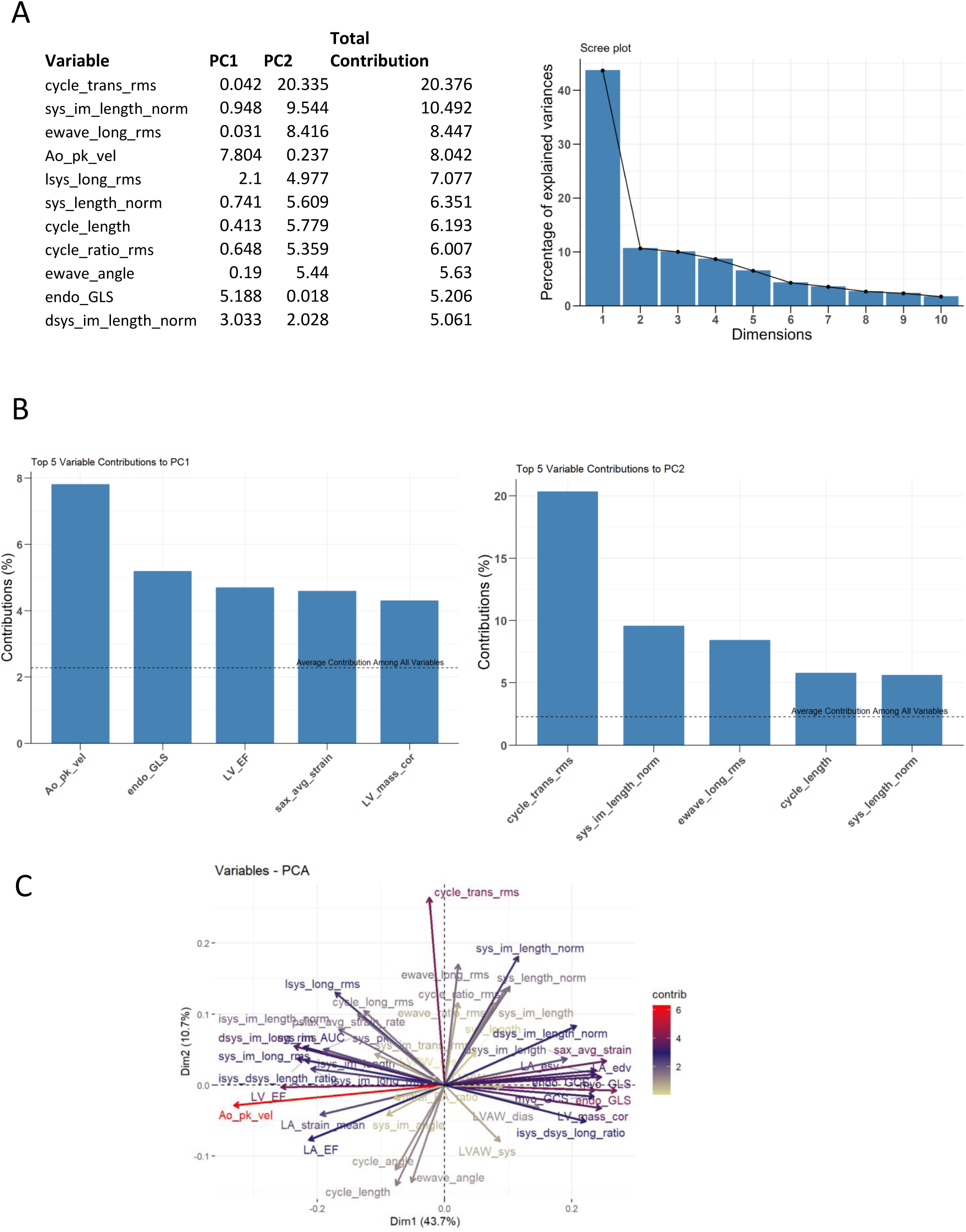
Panels A–C: Principal component analysis showing the variables contributing to the first two principal components.

**Supplemental Table 1.** Traditional cardiac parameter intergroup comparisons in baseline, TAC, and deTAC Groups.

| Metric | Method | Baseline (Mean ± SD) | TAC (Mean ± SD) | deTAC (Mean ± SD) | Baseline vs TAC (p) | Baseline vs deTAC (p) | TAC vs deTAC (p) |
| --- | --- | --- | --- | --- | --- | --- | --- |
| Aortic Peak Velocity | Kruskal-Wallis | -979.86 ± 121.04 | -3737.59 ± 396.37 | -1639.79 ± 311.64 | <0.001 (***) | 0.024 (*) | 0.035 (*) |
| Mitral Valve E-wave to A-wave Ratio | Kruskal-Wallis | 1.89 ± 0.77 | 1.33 ± 0.88 | 1.49 ± 0.60 | 0.033 (*) | 0.267 | 0.228 |
| LV EF | ANOVA | 58.36 ± 4.94 | 37.77 ± 6.54 | 51.52 ± 7.34 | <0.001 (***) | 0.103 | <0.001 (***) |
| Mitral Valve Diameter | ANOVA | 1.64 ± 0.04 | 1.74 ± 0.11 | 1.66 ± 0.17 | 0.192 | 0.942 | 0.323 |
| Aortic Valve Diameter | ANOVA | 1.26 ± 0.04 | 1.38 ± 0.13 | 1.25 ± 0.10 | 0.050 (*) | 0.955 | 0.027 (*) |
| Myocardial GCS | ANOVA | -15.13 ± 1.96 | -9.85 ± 1.92 | -14.87 ± 2.38 | <0.001 (***) | 0.966 | <0.001 (***) |
| Myocardial GLS | ANOVA | -15.46 ± 1.92 | -7.40 ± 2.38 | -11.58 ± 2.79 | <0.001 (***) | 0.01 (*) | 0.006 (**) |
| Endocardial GCS | Kruskal-Wallis | -27.67 ± 3.69 | -17.00 ± 3.34 | -24.93 ± 4.54 | <0.001 (***) | 0.243 | 0.012 (*) |
| Endocardial GLS | ANOVA | -21.46 ± 2.38 | -10.32 ± 2.80 | -16.71 ± 3.62 | <0.001 (***) | 0.012 | <0.001 (***) |
| PLAX Mean Strain Rate | ANOVA | 7.41 ± 1.99 | 5.59 ± 1.83 | 7.56 ± 2.45 | 0.218 | 0.989 | 0.171 |
| SAX Mean Strain | ANOVA | -29.17 ± 2.37 | -16.94 ± 3.73 | -26.29 ± 5.69 | <0.001 (***) | 0.368 | <0.001 (***) |
| LV Normalized Mass | Kruskal-Wallis | 84.09 ± 15.74 | 148.84 ± 11.81 | 105.93 ± 29.61 | <0.001 (***) | 0.157 | 0.035 (*) |
| LV Anterior Wall Diastole | Kruskal-Wallis | 0.88 ± 0.24 | 1.21 ± 0.17 | 0.94 ± 0.14 | 0.01 (*) | 0.724 | 0.015 (*) |
| LV Posterior Wall Diastole | Kruskal-Wallis | 0.77 ± 0.27 | 0.95 ± 0.14 | 0.76 ± 0.06 | 0.013 (*) | 0.416 | 0.063 |
| LV Anterior Wall Systole | Kruskal-Wallis | 1.36 ± 0.30 | 1.54 ± 0.25 | 1.38 ± 0.36 | 0.554 | 0.791 | 0.433 |
| LV Posterior Wall Systole | Kruskal-Wallis | 1.18 ± 0.21 | 1.14 ± 0.17 | 1.12 ± 0.10 | 0.958 | 1 | 1 |

Statistical notations: \* $p < 0.05$ , \*\* $p < 0.01$ , \*\*\* $p < 0.001$ .

**Supplemental Table 2.**
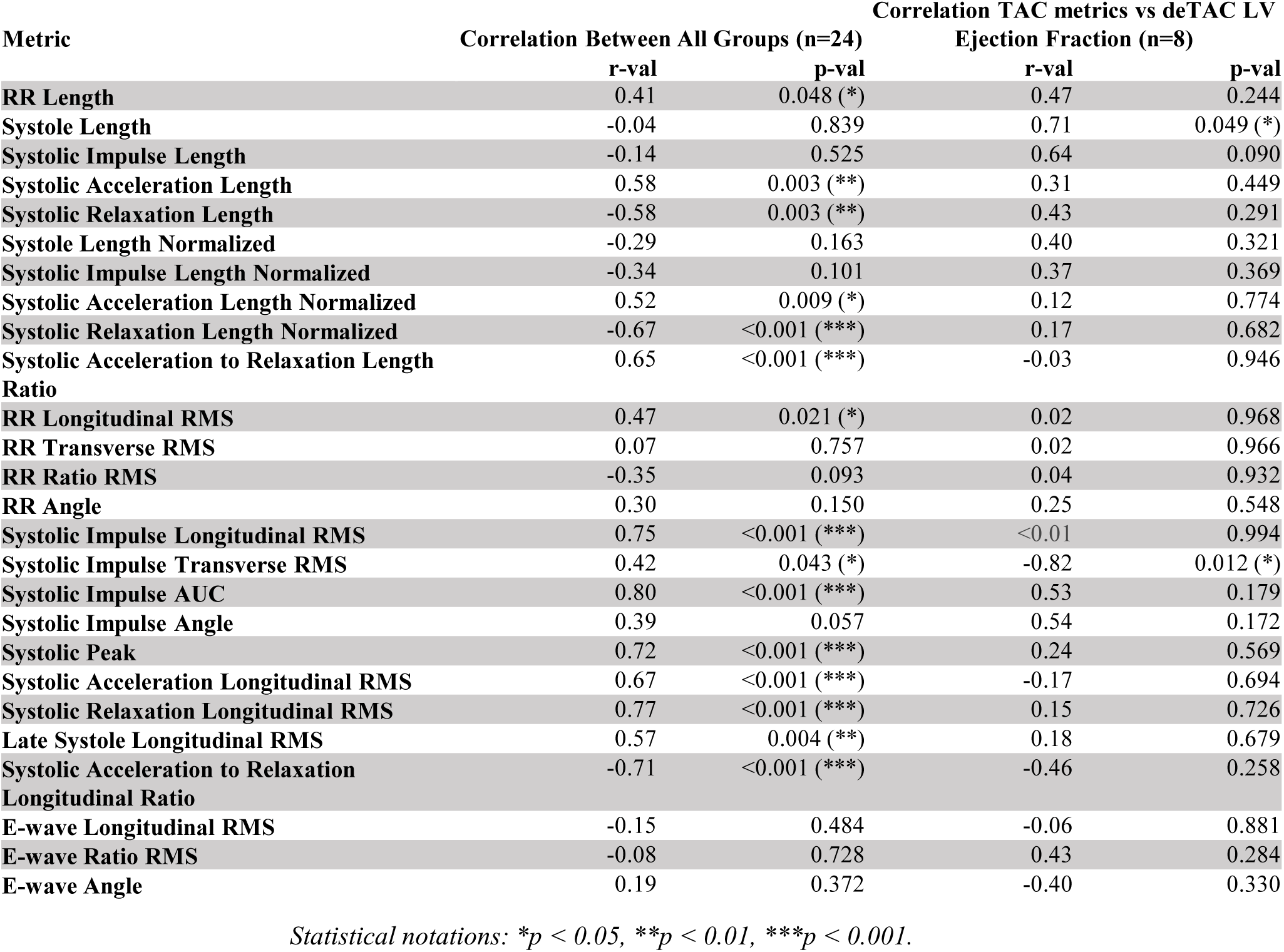
Pearson correlation of HDF cardiac parameters against left ventricular ejection fraction across all conditions and paired-conditions of TAC metrics to deTAC left ventricular ejection fraction.

| Metric | Correlation Between All Groups (n=24) |  | Correlation TAC metrics vs deTAC LV Ejection Fraction (n=8) |  |
| --- | --- | --- | --- | --- |
|  | r-val | p-val | r-val | p-val |
| RR Length | 0.41 | 0.048 (*) | 0.47 | 0.244 |
| Systole Length | -0.04 | 0.839 | 0.71 | 0.049 (*) |
| Systolic Impulse Length | -0.14 | 0.525 | 0.64 | 0.090 |
| Systolic Acceleration Length | 0.58 | 0.003 (**) | 0.31 | 0.449 |
| Systolic Relaxation Length | -0.58 | 0.003 (**) | 0.43 | 0.291 |
| Systole Length Normalized | -0.29 | 0.163 | 0.40 | 0.321 |
| Systolic Impulse Length Normalized | -0.34 | 0.101 | 0.37 | 0.369 |
| Systolic Acceleration Length Normalized | 0.52 | 0.009 (*) | 0.12 | 0.774 |
| Systolic Relaxation Length Normalized | -0.67 | <0.001 (***) | 0.17 | 0.682 |
| Systolic Acceleration to Relaxation Length Ratio | 0.65 | <0.001 (***) | -0.03 | 0.946 |
| RR Longitudinal RMS | 0.47 | 0.021 (*) | 0.02 | 0.968 |
| RR Transverse RMS | 0.07 | 0.757 | 0.02 | 0.966 |
| RR Ratio RMS | -0.35 | 0.093 | 0.04 | 0.932 |
| RR Angle | 0.30 | 0.150 | 0.25 | 0.548 |
| Systolic Impulse Longitudinal RMS | 0.75 | <0.001 (***) | <0.01 | 0.994 |
| Systolic Impulse Transverse RMS | 0.42 | 0.043 (*) | -0.82 | 0.012 (*) |
| Systolic Impulse AUC | 0.80 | <0.001 (***) | 0.53 | 0.179 |
| Systolic Impulse Angle | 0.39 | 0.057 | 0.54 | 0.172 |
| Systolic Peak | 0.72 | <0.001 (***) | 0.24 | 0.569 |
| Systolic Acceleration Longitudinal RMS | 0.67 | <0.001 (***) | -0.17 | 0.694 |
| Systolic Relaxation Longitudinal RMS | 0.77 | <0.001 (***) | 0.15 | 0.726 |
| Late Systole Longitudinal RMS | 0.57 | 0.004 (**) | 0.18 | 0.679 |
| Systolic Acceleration to Relaxation Longitudinal Ratio | -0.71 | <0.001 (***) | -0.46 | 0.258 |
| E-wave Longitudinal RMS | -0.15 | 0.484 | -0.06 | 0.881 |
| E-wave Ratio RMS | -0.08 | 0.728 | 0.43 | 0.284 |
| E-wave Angle | 0.19 | 0.372 | -0.40 | 0.330 |

Statistical notations: \* $p < 0.05$ , \*\* $p < 0.01$ , \*\*\* $p < 0.001$ .

**Supplemental Table 3.** Pearson correlation of traditional cardiac parameters against left ventricular ejection fraction across all conditions and paired-conditions of TAC metrics to deTAC left ventricular ejection fraction.

| Metric | Correlation Between All Groups (n=24) |  | Correlation TAC metrics vs deTAC LV Ejection Fraction (n=8) |  |
| --- | --- | --- | --- | --- |
|  | r-val | p-val | r-val | p-val |
| Aortic Peak Velocity | 0.80 | <0.001 (***) | 0.18 | 0.672 |
| Mitral Valve E-wave to A-wave Ratio | 0.17 | 0.421 | -0.46 | 0.248 |
| LV EF | N/A | N/A | 0.56 | 0.152 |
| Myocardial GCS | -0.93 | <0.001 (***) | -0.65 | 0.084 |
| Myocardial GLS | -0.94 | <0.001 (***) | -0.50 | 0.211 |
| Endocardial GCS | -0.98 | <0.001 (***) | -0.54 | 0.166 |
| Endocardial GLS | -0.94 | <0.001 (***) | -0.49 | 0.214 |
| PLAX Mean Strain Rate | 0.51 | 0.012 (*) | 0.02 | 0.959 |
| SAX Mean Strain | -0.89 | <0.001 (***) | -0.18 | 0.667 |
| LV Normalized Mass | -0.739 | <0.001 (***) | -0.276 | 0.508 |
| LV Anterior Wall Diastole | -0.427 | 0.037 (*) | 0.450 | 0.263 |
| LV Posterior Wall Diastole | -0.322 | 0.125 | -0.050 | 0.907 |
| LV Anterior Wall Systole | -0.191 | 0.372 | 0.422 | 0.297 |
| LV Posterior Wall Systole | 0.229 | 0.283 | 0.234 | 0.577 |

Statistical notations: \* $p < 0.05$ , \*\* $p < 0.01$ , \*\*\* $p < 0.001$ .

